# Habitual foot strike pattern does not affect simulated Triceps Surae muscle metabolic energy consumption during running

**DOI:** 10.1101/779686

**Authors:** Wannes Swinnen, Wouter Hoogkamer, Friedl De Groote, Benedicte Vanwanseele

## Abstract

Foot strike pattern affects ankle joint work and Triceps Surae muscle-tendon dynamics during running. Whether these changes in muscle-tendon dynamics also affect Triceps Surae muscle energy consumption is still unknown. In addition, as the Triceps Surae muscle accounts for a substantial amount of the whole body metabolic energy consumption, changes in Triceps Surae energy consumption may affect whole body metabolic energy consumption. However, direct measurements of muscle metabolic energy consumption during dynamic movements is hard. Model-based approaches can be used to estimate individual muscle and whole body metabolic energy consumption based on Hill type muscle models. In this study, we use an integrated experimental and dynamic optimization approach to compute muscle states (muscle forces, lengths, velocities, excitations and activations) of 10 habitual mid-/forefoot striking and 9 habitual rearfoot striking runners while running at 10 and 14 km/h. The Achilles tendon stiffness of the musculoskeletal model was adapted to fit experimental ultrasound data of the Gastrocnemius medialis muscle during ground contact. Next, we calculated Triceps Surae muscle and whole body metabolic energy consumption using four different metabolic energy models provided in literature. Neither Triceps Surae metabolic energy consumption (p > 0.35), nor whole body metabolic energy consumption (p > 0.14) was different between foot strike patterns, regardless of the energy model used or running speed tested. Our results provide new evidence that mid-/forefoot and rearfoot strike pattern are metabolically equivalent.

## Introduction

The metabolic energy consumed during submaximal running, often referred to as running economy, is an important factor determining endurance running performance (Jones and Carter, 2000). Reduced energy consumption corresponds to improved running economy and hence superior endurance performance (Hoogkamer et al., 2016; Kipp et al., 2019). As such, runners seek to adopt a running pattern with minimal metabolic energy consumption. One aspect of people’s running pattern is foot strike pattern. Although foot strike pattern is a continuum, generally three different foot strike patterns are considered: forefoot strike, midfoot strike and rearfoot strike (Cavanagh and Lafortune, 1980).

While rearfoot striking is the most common running pattern during shod running (Hasegawa et al., 2007; Kasmer et al., 2013; Larson et al., 2011), there seems to be a widespread popular believe that forefoot striking would be more economical than rearfoot striking. Previous research has demonstrated that there is a greater percentage mid-/forefoot strikers among the first finishers in long distance races (de Almeida et al., 2015; Hasegawa et al., 2007), which suggests that forefoot striking may be more economical. However, studies comparing metabolic energy consumption between habitual forefoot and habitual rearfoot strikers found no difference in whole body metabolic energy consumption (Gruber et al., 2013) or even lower energy consumption in rearfoot strikers compared to their forefoot striking colleagues at 11 and 13 km/h but not at 15 km/h (Ogueta-Alday et al., 2014).

Available analyses of the kinetic and kinematic differences between foot strike patterns do not clearly provide evidence for either differences in or unchanged energy consumption with foot strike patterns. The shorter ground contact times (Di Michele and Merni, 2014; Mercer and Horsch, 2015), associated with forefoot striking, may increase metabolic energy consumption according to Kram and Taylor’s cost of generating force hypothesis (Kram and Taylor, 1990). They established that the metabolic energy consumption is inversely proportional to ground contact time, which implies that forefoot strikers may consume more metabolic energy. In addition, forefoot strikers demonstrate greater negative ankle work compared to rearfoot strikers (Stearne et al., 2014). This ankle work is predominantly absorbed by the muscle-tendon unit (MTU) spanning the ankle joint, i.e., Triceps Surae muscle and the in series connected tendinous tissue (SEE, series elastic element). Hence, differences in ankle work may affect the MTU and subsequently the energy consumption of this Triceps Surae muscle. We recently demonstrated that in habitual mid-/forefoot strikers the Gastrocnemius medialis (GM) produces greater muscle force but at lower contraction velocities during early stance compared to habitual rearfoot strikers (Swinnen et al., 2019). Higher muscle force production suggests more muscle activation and thus higher metabolic energy consumption, whereas lower contraction velocities are more force efficient and would therefore reduce muscle activation and thus metabolic energy consumption. Hence, we hypothesized that the differences in metabolic energy consumption would counteract each other and no difference in GM metabolic energy consumption would exist (Swinnen et al., 2019). Yet, as Fletcher and MacIntosh (2017) estimated that 25 to 40% of the total whole body metabolic energy is consumed by the Triceps Surae muscle, we would expect different whole body metabolic energy consumption if Triceps Surae metabolic energy consumption would be different between foot strike patterns.

Model-based approaches have been used to estimate individual muscle and whole body metabolic energy consumption based on Hill type muscle models (Bhargava et al., 2004; Miller, 2014; Uchida et al., 2016; Umberger, 2010; Umberger et al., 2003). However, to obtain reliable simulation results, a close match between simulated and experimental data is essential. Here, we used experimental dynamics ultrasound data from the Gastrocnemius medialis (GM) to improve our dynamic optimization and as such, ensure more reliable estimations of muscle metabolic energy consumption. We used four different metabolic energy models (Bhargava et al., 2004; Uchida et al., 2016; Umberger, 2010; Umberger et al., 2003) to calculate Triceps Surae muscle and whole body metabolic energy consumption of habitual mid-/forefoot and rearfoot strikers running at 10 and 14 km/h. We hypothesized that neither Triceps Surae nor whole body metabolic energy consumption would be different between foot strike patterns.

## Methods

### Participants

Ten habitual mid-/forefoot strikers (6 males, 4 females; body mass: 65.2 ± 7.7 kg; body height: 1.78 ± 0.07 m) and 9 habitual rearfoot strikers (6 males, 3 females; body mass: 72.7 ± 12.5 kg; body height: 1.81 ± 0.08 m) participated in this study. All participants were regular runners who ran at least 30 km/week, did not have any Achilles tendon or calf injury in the last six months and had no prior Achilles tendon surgery. Written informed consent, approved by the local ethical committee (Medical Ethical Committee of UZ Leuven), was obtained at the start of the experiment.

### Experimental procedure

The experimental procedures have been described in detail in our earlier publication on gastrocnemius medialis muscle-tendon interaction and muscle force production in this group of runners (Swinnen et al., 2019). Briefly, after a 10 minutes warm-up, participants ran 5 minutes on a force measuring treadmill (Motekforce Link, Amsterdam, The Netherlands): 2.5 minutes at 10 and at 14 km/h, in randomized order. We collected kinetic, kinematic, muscle activation and ultrasound data of at least four strides during the last minute of each running speed. All measurements were synchronized through a trigger pulse signal sent from the ultrasound device.

### Kinetics, kinematic and foot strike angle

Thirteen infrared cameras (Vicon, Oxford Metrics, UK) captured the motion of forty-seven reflective markers at a sampling frequency of 150 Hz. We used OpenSim 3.3 (OpenSim, Stanford, USA) to first scale a musculoskeletal model based on the subject’s dimensions (Hamner et al., 2010) and to subsequently compute joint kinematics using a Kalman smoothing algorithm (De Groote et al., 2008). Muscle tendon unit lengths were calculated using OpenSim’s Muscle Analysis Tool.

Ground reaction force data, sampled at 900 Hz, was first low pass filtered with a cut-off frequency of 20 Hz and used to determine ground contact phase adopting a 30 N threshold. We determined foot strike angle using a marker based method (Altman and Davis, 2012). At initial ground contact, we drew a line through the first metatarsal-phalangeal joint marker and heel marker of the left foot. The angle between this line and the ground was calculated and considered as the foot strike angle. Following Altman and Davis (2012) runners with a foot strike angle greater than 8° were considered rearfoot strikers, while runners with a foot strike angle under 8° were considered mid-/forefoot strikers. Foot strike angle was averaged over the strides used for ultrasound analysis. Foot strike type (rearfoot or mid-/forefoot) was consistent within subjects across running speeds.

We calculated joint torques using OpenSim’s Inverse Dynamics Tool based on joint kinematics and ground reaction forces. Joint torques were low pass filtered using a recursive fourth order Butterworth filter with cut-off frequency of 20 Hz.

### Dynamic ultrasound imaging

We collected dynamic ultrasound images of the GM muscle fascicles of the left leg with a B-mode ultrasound system (Telemed Echoblaster 128 CEXT system) sampling at 86 Hz. The linear transducer (UAB Telemed, Vilnius, Lithuania, LV 7.5/60/128Z-2) was placed on the mid-belly of the muscle, aligned with the muscle fascicles and attached to the calf with tape and bandages. To analyze the GM muscle fascicle lengths and pennation angles we used a semiautomatic tracking algorithm (Farris and Lichtwark, 2016). We analyzed at least four strides and calculated fascicle length changes relative to fascicle length at toe-off. All data were splined to 100 data points per ground contact, starting at initial contact.

### Muscle activity

We used surface electromyography (EMG) to determine GM and Soleus (SOL) muscle activity of the right leg through a wireless EMG acquisition system (ZeroWire EMG Aurion, Milano, Italy) measuring at 900 Hz. EMG signals were first band-pass filtered (20-400 Hz), rectified and low-pass filtered (20 Hz). For each subject and muscle, EMG waveforms were normalized to maximal activation, determined as the maximal activation of each muscle using a moving average over 10 data points. Due to technical issues, the EMG data of the GM of one participant (mid-/forefoot striker) and SOL of three participants (2 mid-/forefoot strikers and 1 rearfoot striker) could not be used.

Comparison between experimental EMG and simulated activation of the GM and SOL demonstrated similar trends, yet due to our optimization criteria (minimization of muscle activation squared) preactivation is not predicted (Fig. S1).

### Estimating muscle and whole body metabolic energy consumption

Several models for estimating muscle metabolic energy rate have been proposed and it is yet unclear which model yields the most valid results. We, therefore, used multiple models primarily to assure that our results are independent from the metabolic energy model used. Our goal was not to compare the different energy models (for comparison between metabolic energy models see Miller 2014). All models required the muscle states (i.e., muscle activations, excitations, lengths, velocities and forces) as inputs. To obtain these muscle states we solved the muscle redundancy problem using a dynamic optimization algorithm that takes into account muscle-tendon dynamics (i.e., muscle activation and contraction dynamics) of the 43 lower limb muscles of the left leg in our model (De Groote et al., 2009; De Groote et al., 2016). Individual muscle moment arms, muscle tendon unit lengths and muscle properties were extracted from the scaled OpenSim model and were input to the muscle redundancy solver. We scaled maximal isometric muscle force based on the subject’s body mass and height (Handsfield et al., 2014). To avoid maximal muscle activations and unrealistically high reserve actuator forces, muscle forces were multiplied by 3 for all participants. The Triceps Surae muscles, containing the GM, Gastrocnemius lateralis (GL) and SOL, were modeled as three separate muscle-tendon units, with the tendon representing the Achilles tendon. To ensure a close match between experimental GM muscle fascicle length changes and simulated GM muscle fascicle lengths, we adjusted the normalized tendon stiffness, a scaling factor to calculate GM, GL, SOL tendon stiffness based on the ratio between maximal isometric force and tendon slack length, to a value of 5 for all participants (Figure 1). Gerus et al. (2015) previously stated that the Achilles tendon is more compliant than the generic tendon stiffness as described by Zajac (1989). We tested multiple other values (ranging from 4 to 35) where 5 gave the best match. The normalized stiffness for all other muscles was kept on the default value of 35. Joint torques served as inputs to solve the muscle redundancy problem by minimizing the squared muscle activation. We solved the dynamic optimization problem through direct collocation using GPOP-II software (Patterson and Rao, 2014). Subsequently the resulting nonlinear equations was solved using ipopt (Wächter and Biegler, 2006). In 9 out of the 154 ground contact analyzed the optimization algorithm failed to find an optimal solution, these strides were excluded.

**Figure 1.**
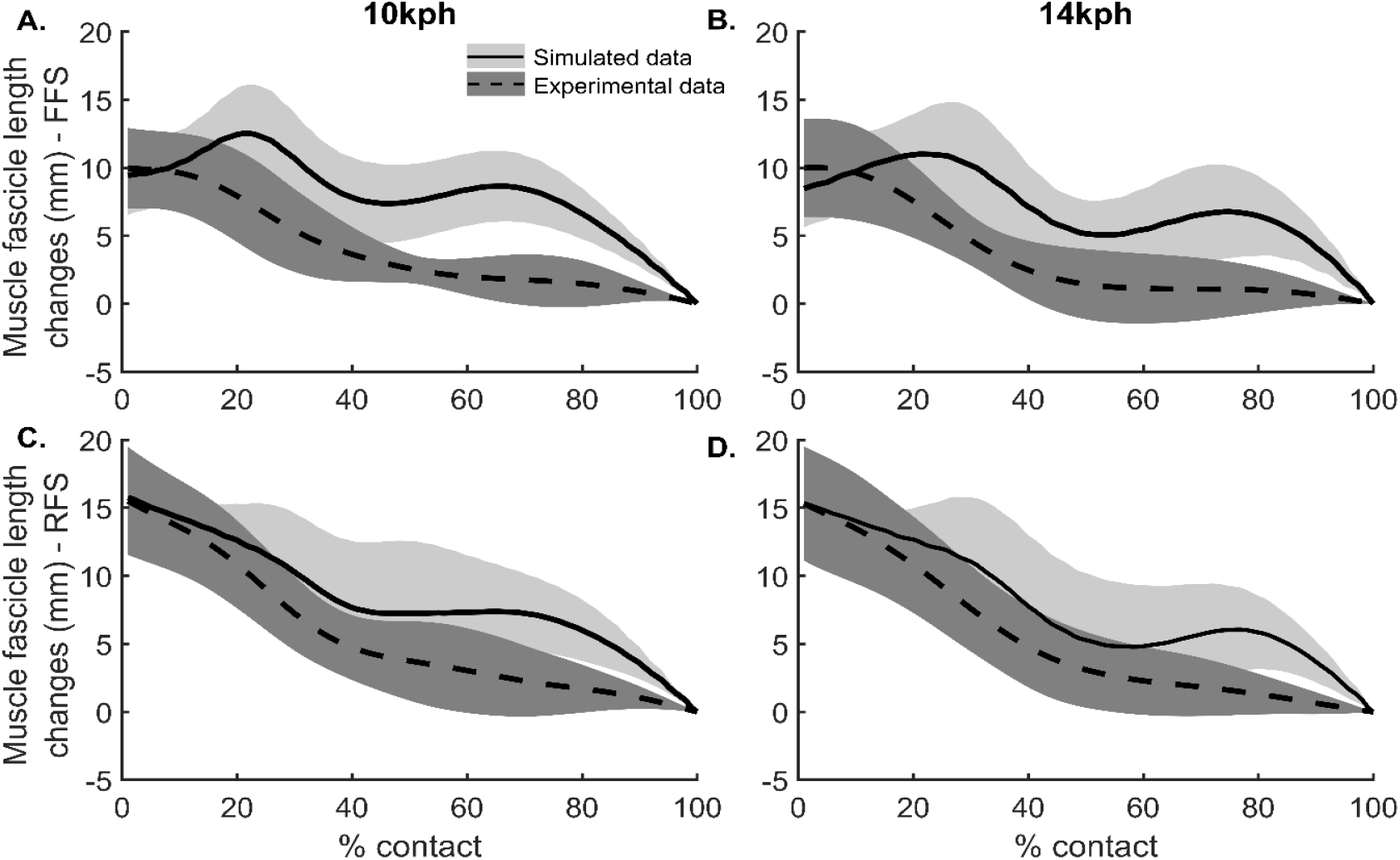
Simulated (solid) and experimental (dashed) GM muscle fascicle length changes during ground contact in mid-/forefoot strikers (A,B; n = 10) and rearfoot strikers (C,D; n = 9) at 10 km/h (A,C) and 14 km/h (B,D). Muscle fascicle length changes are normalized to muscle fascicle length at toe off. Shaded area represent standard deviation.

Next, the simulated muscle states were used as input in four models to estimate muscle metabolic energy rate 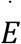 that are consistent with Hill based muscle dynamics: Umberger, Gerritsen and Martin (2003) (U2003), Bhargava, Pandy and Anderson (2004) (B2004), Umberger (2010) (U2010) and Uchida *et al*. (2016) (U2016). All these models had the same general form to calculate energy expenditure:

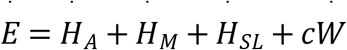

Where *Ḣ_A_, Ḣ_M_* and *Ḣ_SL_* are the heat production rates of the muscles for activation, maintenance and shortening/lengthening respectively, *Ẇ* is the muscle mechanical work rate where concentric work is defined positively and *C* is weighting factor depending on the type of work (concentric of eccentric). Major differences between the models are how they treat eccentric muscle work and how they weight muscle lengthening heat rate. While in U03 and U16 negative mechanical work (i.e., metabolic energy generation) is incorporated, B04 and U10 are restricted to positive mechanical work only, negative mechanical work is excluded and the lengthening heat rate coefficient is adapted. Apart from these differences, the heat rate calculations have similar terms between the models, though the scaling factors used are different. Activation and maintenance heat rates are generally defined by muscle mass/force, length and fiber type composition while shortening/lengthening heat rate depend on muscle contraction velocity. U03, U10 and U16 scale these heat rates by muscle activation whereas B04 does not. We refer to the specific papers for more detailed information on the models.

Muscle metabolic energy rate was integrated over time to obtain metabolic energy consumption during one stance phase which was then multiplied by 2, to account for both legs, and multiplied by the stride frequency to obtain metabolic energy rate in Watts. The metabolic energy consumed by the Triceps Surae muscles was normalized to their respective muscle mass. We computed whole body metabolic energy expenditure as the sum of metabolic energy consumed by all 43 muscles included in the model and added a basal rate of 1.2 W/kg (Waters and Mulroy, 1999). Whole body metabolic energy consumption was normalized to body mass.

### Statistics

All data are presented as mean ± standard deviation. We categorized our data in four groups: mid-/forefoot strike at 10 km/h (FF 10), mid-/forefoot strike at 14 km/h (FF 14), rearfoot strike at 10 km/h (RF 10) and rearfoot strike at 14 km/h (RF 14). First, normality was checked with the Shaprio-Wilk test. If data from all groups followed a normal distribution a mixed analysis of variance (ANOVA) was used to determine interaction and main effects (foot strike pattern and running speed) using SPSS v.24 (IBM SPSS, Armonk, New York, USA). Yet, if not all the data in the groups followed a normal distribution, the non-parametric Mann-Whitney U test was performed to compare foot strike pattern differences at 10 and 14 km/h separately. To determine the effect of running speed for these datasets, the data was first grouped according to running speed and again checked upon normality. If both datasets were then normally distributed, a paired t-test was performed, if not we performed a Wilcoxon signed-rank test. Statistical significance was considered when p < 0.05.

## Results

Although mean foot strike angle was more than 15° different between both foot strike groups (p < 0.01; Table 1), Triceps Surae metabolic energy consumption was not different between foot strike patterns, regardless of speed or metabolic energy model (p > 0.35; Figure 2). Moreover, metabolic energy consumed by the individual Triceps Surae muscles, i.e. GM, GL and SOL, was not different between foot strike patterns (p > 0.10) independent of the model used or running speed. Furthermore, estimated whole body metabolic energy consumption was not different between foot strike patterns regardless of model or running speed tested (p > 0.14; Figure 3). As one would expect, running faster resulted in greater metabolic energy consumption in the Triceps Surae muscle group (p < 0.01) as well as in all three Triceps Surae individually (p < 0.02). Also, whole body metabolic energy consumption was greater when running at 14 km/h compared to 10 km/h (p < 0.01).

**Figure 2.**
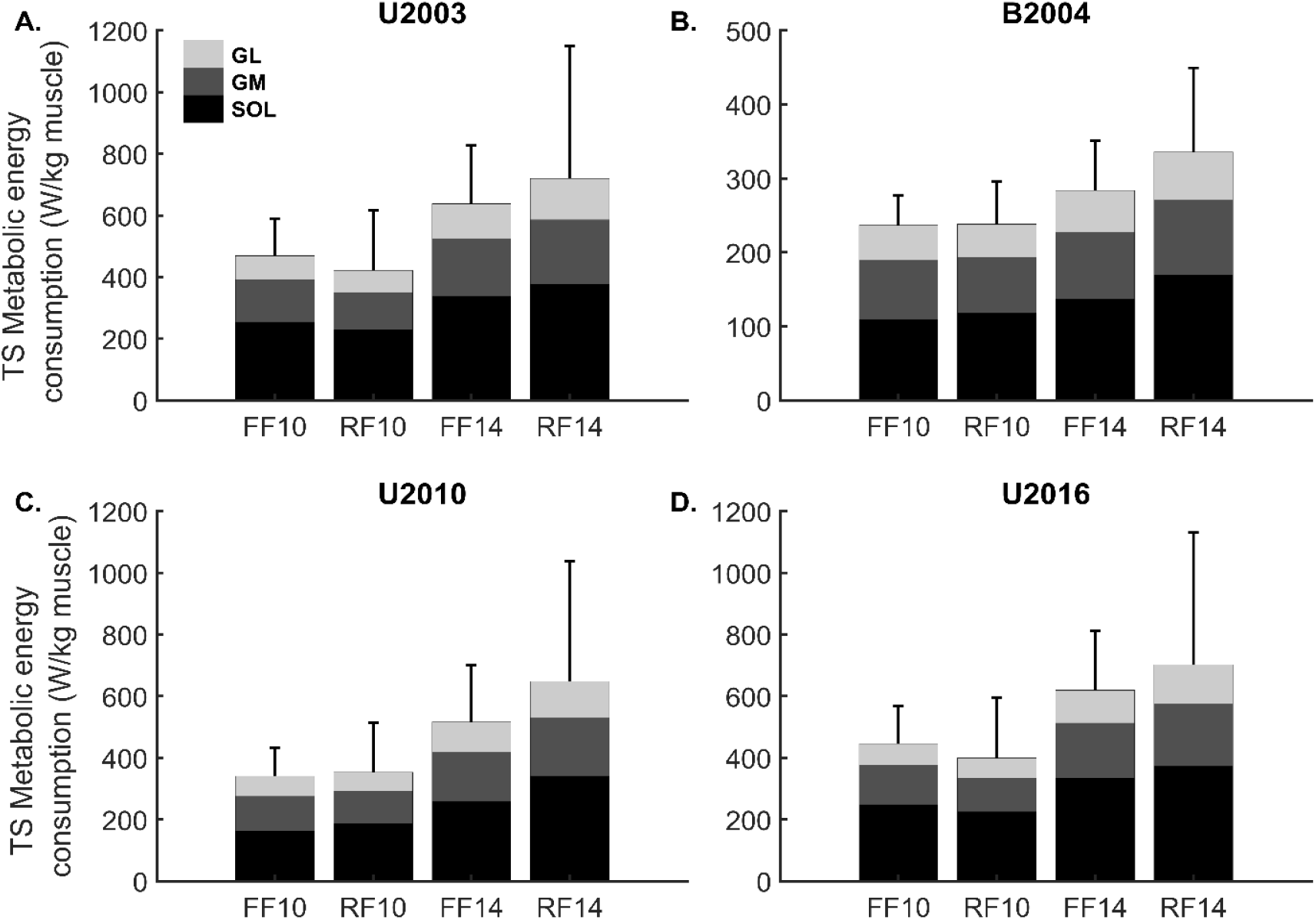
Triceps Surae (TS) metabolic energy consumption including individual muscles: Soleus (black), Gastrocnemius medialis (dark grey) and Gastrocnemius lateralis (light grey) in mid-/forefoot strikers (FF, n = 10) and rearfoot strikers (RF, n = 9). Mixed ANOVA or Mann-Whitney U test demonstrated no significant difference in metabolic energy consumed between foot strike patterns, not for individual Triceps Surae muscle (p > 0.10) nor for all three muscles together (p > 0.35). Mixed ANOVA, paired t-test or Wilcoxon signed-rank test demonstrated significant greater energy consumption at 14km/h compared to 10km/h (p < 0.01).

**Table 1.** Comparison between mid-/forefoot and rearfoot strikers and between 10 and 14 km/h. All data are expressed as mean ± SD. ^a^ significant main foot strike effect. ^b^ significant running speed effect.^c^significant interaction effect.

|  |  | speed | Forefoot strike | Rearfoot strike |
| --- | --- | --- | --- | --- |
| Foot strike angle<br>(°)<sup>a</sup> | | 10 km/h | -0.4 ± 4.4 | 14.8 ± 3.7 |
| | | 14 km/h | 0.3 ± 5.3 | 17.2 ± 5.4 |
| Ratio (%)<br>($E_{TS}/E_{WB}$) | U03<sup>c</sup> | 10 km/h | 26 ± 4 | 22 ± 8 |
| | | 14 km/h | 25 ± 3 | 25 ± 8 |
| | B04<sup>b</sup> | 10 km/h | 26 ± 3 | 27 ± 6 |
| | | 14 km/h | 23 ± 4 | 26 ± 6 |
| | U10<sup>b</sup> | 10 km/h | 27 ± 4 | 28 ± 9 |
| | | 14 km/h | 28 ± 5 | 32 ± 10 |
| | U16<sup>c</sup> | 10 km/h | 27 ± 4 | 23 ± 8 |
| | | 14 km/h | 26 ± 3 | 26 ± 9 |

The ratio of metabolic energy consumed by the Triceps Surae relative to whole body metabolic energy consumption ranged between 22 and 32% across foot strike patterns and running speeds but was not different between foot strike patterns (p > 0.19). In contrast, the different models revealed inconsistent results when the effect of speed on this ratio was considered. While U03 and U16 did not show significant differences in this ratio between running speeds (p > 0.07), U10 showed a significant greater ratio at 14km/h compared to 10 km/h (p = 0.01), whereas B04 showed a significant smaller ratio at 14 km/h than at 10 km/h (p = 0.02).

## Discussion

This study investigated the effect of habitual foot strike pattern on simulated Triceps Surae muscle and whole body metabolic energy consumption. We used a dynamic optimization approach in which the Achilles tendon stiffness of the musculoskeletal model was adapted to better match experimental GM ultrasound data (Figure 1). Four different metabolic energy models were incorporated to ensure model independency. In line with our hypothesis, none of the individual Triceps Surae muscles, nor whole body metabolic energy consumption demonstrated significant differences between mid-/forefoot strikers and rearfoot strikers (Figure 2 and Figure 3). Faster running increased both simulated Triceps Surae muscle and whole body metabolic energy consumption.

**Figure 3.**
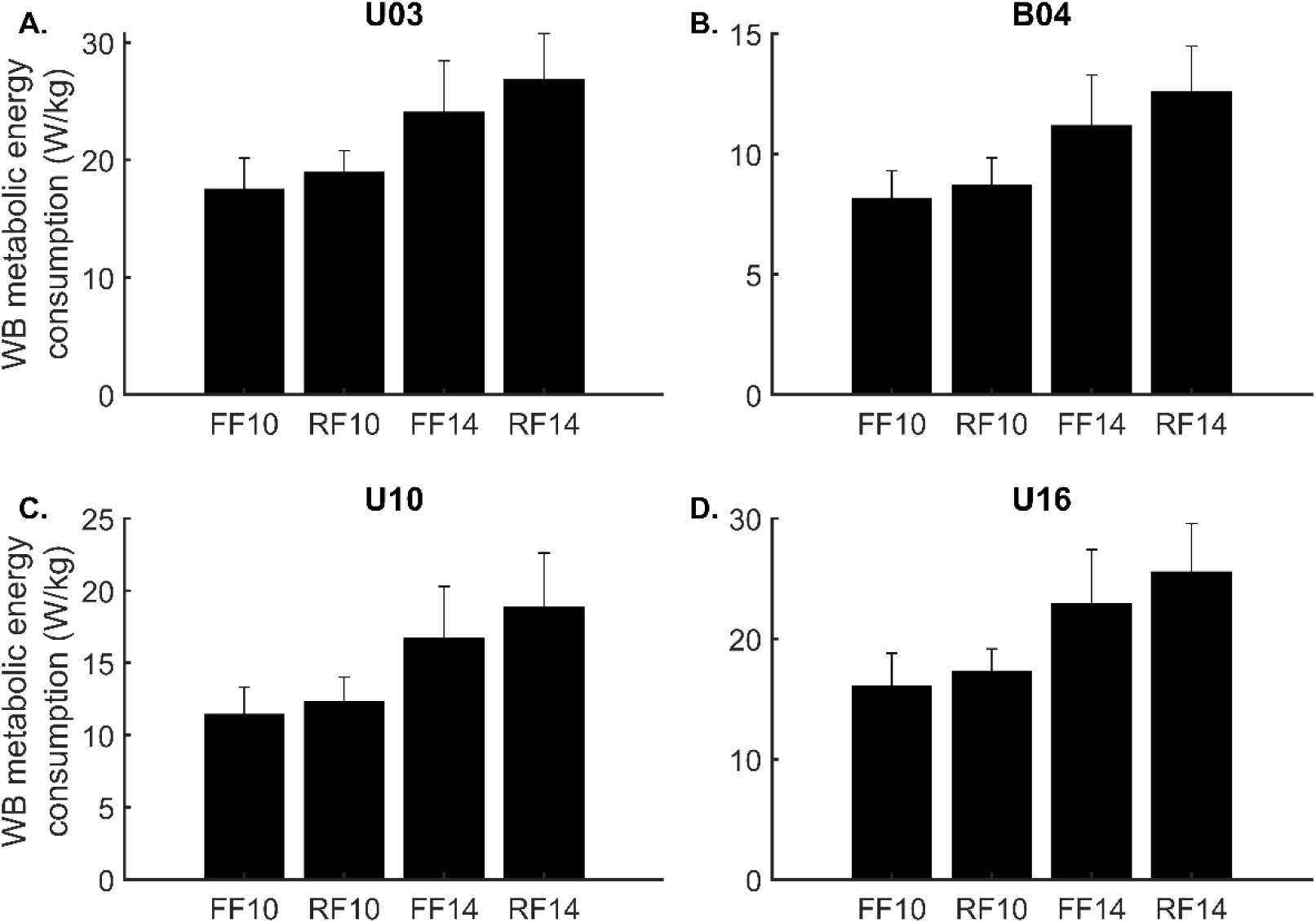
Estimated whole body (WB) metabolic energy consumption for all four metabolic energy models used for mid-/forefoot strikers at 10 km/h (FF10) and 14 km/h (FF14) and rearfoot strikers at 10 km/h (RF10) and 14 km/h (RF14). U03 = Umberger, Gerritsen and Martin (2003), B04 = Bhargava, Pandy and Anderson (2004), U10= Umberger (2010) and U16 = Uchida et al. (2016). Mixed ANOVA or Mann-Whitney U test demonstrated no significant difference between foot strike patterns (p > 0.14). Mixed ANOVA, paired t-test or Wilcoxon signed-rank test demonstrated significant increase in energy consumption when running at 14 km/h compared to 10 km/h.

Our results provide additional scientific evidence that mid-/forefoot and rearfoot strike patterns are energetically equivalent. We recently showed that GM muscle force production is greater while muscle contraction velocity is smaller in mid-/forefoot strikers compared to rearfoot strikers, especially during early ground contact (Swinnen et al., 2019). Here, we provide further evidence that the greater muscle forces in mid-/forefoot strikers are more economically produced due to the lower muscle contraction velocities and hence no difference in GM, GL or SOL metabolic energy consumption between foot strike patterns exist. Moreover, previous experimental research already demonstrated that differences in whole body metabolic energy consumption between foot strike patterns are small (Ogueta-Alday et al., 2014) or even non-existing (Cunningham et al., 2010; Gruber et al., 2013; Lussiana et al., 2017; Perl et al., 2012). Studies investigating the effect of gait retraining from rearfoot to forefoot strike running do not find an effect on the metabolic energy consumption during running when enough training sessions (?8) were offered (Ekizos et al., 2018; Roper et al., 2017). However, when only two training sessions were provided an initial increase in metabolic cost is reported (Ekizos et al., 2018), indicating the need for habituation. Hence, since habituation is necessary when switching foot strike pattern and switching ultimately does not result in a reduced metabolic cost, switching foot strike pattern seems to be ineffective from a performance point of view.

Next to estimated Triceps Surae muscle and whole body metabolic energy rate, the contribution of the Triceps Surae to the whole body metabolic energy rate (i.e. ratio) was also not different between foot strike patterns. However, the effect of running speed was less clear. Two models (U03 and U16) did not find a speed effect, while U10 and B04 did find a speed effect, but in opposing directions. With faster running the relative contribution of joint power/work during ground contact seems to gradually shift more towards proximal joints (i.e. hip), especially at running speeds closer to sprinting (Schache et al., 2015). Hence, if a shift in muscle metabolic energy consumption would occur, a shift in the same direction as joint power would have been expected, implying a decreased relative contribution of the Triceps Surae with increasing running speed. However, the difference in running speeds tested in this study was small and our fastest speed did not approach sprinting. Therefore, to better understand the effect of running speed on the distribution of muscle metabolic energy consumption across lower extremity muscles a wider range of running speeds should be investigated.

Dynamic optimization allowed us to account for muscle-tendon interactions when estimating muscle states. A good match between experimental and predicted muscle states is crucial for good estimations of muscle metabolic energy. We found that it was important to adapt Achilles tendon stiffness to obtain a close match between simulated and measured GM fiber lengths. Using a generic normalized tendon stiffness value of 35 resulted in negligible length changes of the tendinous tissues and as a consequence muscle fascicle length changes were no longer uncoupled from length changes of the entire muscle tendon unit (Fig. S2). Nevertheless, there is ample experimental evidence that the tendinous tissue interacts with the Triceps Surae muscles, uncoupling the muscle fascicle length changes from the length changes of the entire MTU (Fukunaga et al., 2002; Lai et al., 2015; Lichtwark and Wilson, 2008), allowing the muscle fascicles to contract at much slower - more force-efficient - velocities implying lower metabolic energy consumption (Hill, 1922; van der Zee, Lemaire and van Soest, 2019). As a result, predicted Triceps Surae muscle metabolic energy consumption with the generic stiff tendon was on average 80% higher compared to the adapted Achilles tendon stiffness (Fig. S3). Also, estimated whole body metabolic energy consumption was on average 23% higher compared to the adapted Achilles tendon stiffness (Fig. S4). The discrepancy between the results based on the generic and adapted tendon stiffness values illustrates the importance of a good match between computed and experimental muscle states to obtain reliable results of muscle metabolic energy consumption. Moreover, the increased metabolic energy consumption associated with the stiff tendon emphasizes the importance of the muscle-tendon unit interaction on the metabolic energy consumption during running.

Although our conclusions are independent of the metabolic energy model used, the wide variability in absolute energy rates between the metabolic energy models are remarkable. While B04 and U10 predict experimental whole body metabolic energy consumption rather close to experimental data, whole body metabolic energy consumption predicted by U03 and U16 are almost twice as high as experimentally observed (Batliner et al., 2018; Kipp et al., 2018). The major difference is that U03/U16 neglect eccentric work whereas B04/U10 account for eccentric work. Instead of accounting for negative work, U03/U16 reduce the lengthening heat rate coefficient. Our results (lower energy rates with U03/U16) illustrate that the reduction of the lengthening heat rate more than offsets the exclusion of eccentric muscle work. While we seem to have a good understanding of the energy cost of isometric and concentric muscle contractions, the energy cost during eccentric or stretch-shortening muscle contraction is more debatable. It is clear that eccentric muscle work is more efficiently produced compared to concentric muscle work (Hill, 1960), and therefore it appears reasonable to allow eccentric muscle work and muscle lengthening to reduce the metabolic energy consumption rate of a muscle, however a clear consensus on how to treat eccentric work is still lacking. Also the energy cost associated with the stretch-shortening of a muscle is still controversial (Holt et al., 2014; van der Zee et al., 2019). Nevertheless, in contrast to the absolute differences, the relative increase in metabolic energy consumption based on all muscle metabolic models when running faster corresponds quite well with the experimental data. Experimental data indicates that increasing the running speed from 10 km/h to 14 km/h would corresponds with an increase in whole body metabolic energy consumption of approximately 40 to 45% (Batliner et al., 2018; Kipp et al., 2018). The energy models predict similar increases of 40% (U03), 41% (B04), 49% (U10) and 45% (U16). In summary, while metabolic energy models do a good job for predicting relative changes, absolute values are not in accordance with experimental data. Therefore, experimental muscle research on how to account for the energy cost of eccentric and stretch-shortening muscle contractions is necessary before recommendations on how to implement these contractions in metabolic energy models can be made.

Our study has some limitations. First, we did not measure Achilles tendon stiffness from our participants and assumed equal normalized Achilles tendon stiffness for all subjects. Kubo *et al*. (2015) found no difference in Achilles tendon stiffness between foot strike patterns and thus, on average, we can assume equal normalized Achilles tendon stiffness. Mid-/forefoot strikers are reported to earlier activate their Gastrocnemii muscles (Ahn et al., 2014; Swinnen et al., 2019). However due to our optimization criteria (i.e. minimization of muscle activation squared) pre-activation of the Triceps Surae muscles is not predicted. Still, our simulations demonstrate a slightly earlier Triceps Surae muscle activation in mid-/forefoot strikers than rearfoot strikers (Fig. S1). Furthermore, our musculoskeletal model has some limitations. For example, the musculoskeletal model lacks a midfoot arch, which has been shown to store and release energy and subsequently reduce the metabolic rate during running (Ker et al., 1987; Stearne et al., 2016). Moreover, we only took metabolic energy expenditure during ground contact into account, according to Arellano and Kram (2014) only considering ground contact would lead to an underestimation of 7% of the net metabolic energy expenditure. We used ultrasound data to validate our simulations, a well-known limitation of ultrasound data is that these 2D images represents a 3D muscle structure, possibly resulting in underestimation of muscle fascicle length changes when there is out of plane muscle movement.

In conclusion, we demonstrated that – in contrast with the widespread belief in the running community – none of the foot strike patterns induce a reduction in metabolic energy consumption of the Triceps Surae muscle while running. In agreement with previous experimental research, simulated whole body metabolic energy consumption was also similar between foot strike patterns. Hence, we conclude that none of the foot strike patterns can be associated with a superior running energetics. Yet, we looked into differences in metabolic rate during sub-maximal running, an important performance parameter in distance running. It should be noted that for sprinting energy rate is not as important due to the short distance/time.

## References

Ahn, A. N., Brayton, C., Bhatia, T. and Martin, P. (2014). Muscle activity and kinematics of forefoot and rearfoot strike runners. J. Sport Heal. Sci. 3, 102–112.

Altman, A. R. and Davis, I. S. (2012). A kinematic method for footstrike pattern detection in barefoot and shod runners. Gait Posture 35, 298–300.

Arellano, C. J. and Kram, R. (2014). Partitioning the metabolic cost of human running: A task-by-task approach. Integr. Comp. Biol. 54, 1084–1098.

Batliner, M. E., Kipp, S., Grabowski, A. M., Kram, R., Byrnes, W. C., Physiology, I. and States, U. (2018). Does Metabolic Rate Increase Linearly with Running Speed in all Distance Runners? Sport. Med. Int. Open 2, E1–E8.

Bhargava, L. J., Pandy, M. G. and Anderson, F. C. (2004). A phenomenological model for estimating metabolic energy consumption in muscle contraction. J. Biomech. 37, 81–88.

Cavanagh, P. R. and Lafortune, M. A. (1980). Ground reaction forces in distance running. J. Biomech. 13, 397–406.

Cunningham, C. B., Schilling, N., Anders, C. and Carrier, D. R. (2010). The influence of foot posture on the cost of transport in humans. J. Exp. Biol. 213, 790–797.

de Almeida, M. O., Saragiotto, B. T., Yamato, T. P. and Lopes, A. D. (2015). Is the rearfoot pattern the most frequently foot strike pattern among recreational shod distance runners? Phys. Ther. Sport 16, 29–33.

De Groote, F., De Laet, T., Jonkers, I. and De Schutter, J. (2008). Kalman smoothing improves the estimation of joint kinematics and kinetics in marker-based human gait analysis. J. Biomech. 41, 3390–3398.

De Groote, F., Pipeleers, G., Jonkers, I., Demeulenaere, B., Patten, C., Swevers, J. and De Schutter, J. (2009). A physiology based inverse dynamic analysis of human gait: Potential and perspectives. Comput. Methods Biomech. Biomed. Engin. 12, 563–574.

De Groote, F., Kinney, A. L., Rao, A. V. and Fregly, B. J. (2016). Evaluation of Direct Collocation Optimal Control Problem Formulations for Solving the Muscle Redundancy Problem. Ann. Biomed. Eng. 44, 2922–2936.

Di Michele, R. and Merni, F. (2014). The concurrent effects of strike pattern and ground-contact time on running economy. J. Sci. Med. Sport 17, 414–418.

Ekizos, A., Santuz, A. and Arampatzis, A. (2018). Short- and long-term effects of altered point of ground reaction force application on human running energetics. J. Exp. Biol. 221,.

Farris, D. J. and Lichtwark, G. A. (2016). UltraTrack: Software for semi-automated tracking of muscle fascicles in sequences of B-mode ultrasound images. Comput. Methods Programs Biomed. 128, 111–118.

Fletcher, J. R. and MacIntosh, B. R. (2017). Running economy from a muscle energetics perspective. Front. Physiol. 8,.

Fukunaga, T., Kawakami, Y., Kubo, K. and Kanehisa, H. (2002). Muscle and Tendon Interaction During Human Movements. Exerc. Sport Sci. Rev. 30, 106–110.

Gerus, P., Rao, G. and Berton, E. (2015). Ultrasound-based subject-specific parameters improve fascicle behaviour estimation in Hill-type muscle model. Comput. Methods Biomech. Biomed. Engin. 18, 116–123.

Gruber, A. H., Umberger, B. R., Braun, B. and Hamill, J. (2013). Economy and rate of carbohydrate oxidation during running with rearfoot and forefoot strike patterns. J. Appl. Physiol. 115, 194–201.

Hamner, S. R., Seth, A. and Delp, S. L. (2010). Muscle contributions to propulsion and support during running. J. Biomech. 43, 2709–2716.

Handsfield, G. G., Meyer, C. H., Hart, J. M., Abel, M. F. and Blemker, S. S. (2014). Relationships of 35 lower limb muscles to height and body mass quantified using MRI. J. Biomech. 47, 631–638.

Hasegawa, H., Yamauchi, T. and Kraemer, W. J. (2007). Foot strike patterns of runners at the 15-km point during an elite-level half marathon. J. Strength Cond. Res. 21, 888–893.

Hill, A. V (1922). The maximum work and mechanical efficiency of human muscles, and their most economical speed. J. Physiol. 56, 19–41.

Hill, A. V. (1960). Production and absorption of work by muscle. Science 131, 897–903.

Holt, N. C., Roberts, T. J. and Askew, G. N. (2014). The energetic benefits of tendon springs in running: is the reduction of muscle work important? J. Exp. Biol. 217, 4365–4371.

Hoogkamer, W., Kipp, S., Spiering, B. A. and Kram, R. (2016). Altered running economy directly translates to altered distance-running performance. Med. Sci. Sports Exerc. 48, 2175–2180.

Jones, A. M. and Carter, H. (2000). The effect of endurance training on parameters of aerobic fitness. Sport. Med. 29, 373–386.

Kasmer, M. E., Liu, X. C., Roberts, K. G. and Valadao, J. M. (2013). Foot-strike pattern and performance in a marathon. Int. J. Sports Physiol. Perform. 8, 286–292.

Ker, R. F., Bennett, M. B., Bibby, S. R., Kester, R. C. and Alexander, R. M. (1987). The spring in the arch of the human foot. Nature 325, 147–149.

Kipp, S., Grabowski, A. M. and Kram, R. (2018). What determines the metabolic cost of human running across a wide range of velocities? J. Exp. Biol. 221, jeb184218.

Kipp, S., Kram, R. and Hoogkamer, W. (2019). Extrapolating metabolic savings in running: Implications for performance predictions. Front. Physiol. 10,.

Kram, R. and Taylor, R. C. (1990). Energetics of running: a new perspective. Nature 346, 265–267.

Kubo, K., Miyazaki, D., Tanaka, S., Shimoju, S. and Tsunoda, N. (2015). Relationship between Achilles tendon properties and foot strike patterns in long-distance runners. J. Sports Sci. 33, 665–669.

Lai, A., Lichtwark, G. A., Schache, A. G., Lin, Y.-C., Brown, N. A. T. and Pandy, M. G. (2015). In vivo behavior of the human soleus muscle with increasing walking and running speeds. J. Appl. Physiol. 118, 1266–1275.

Larson, P., Higgins, E., Kaminski, J., Decker, T., Preble, J., Lyons, D., McIntyre, K. and Normile, A. (2011). Foot strike patterns of recreational and sub-elite runners in a long-distance road race. J. Sports Sci. 29, 1665–1673.

Lichtwark, G. A. and Wilson, A. M. (2008). Optimal muscle fascicle length and tendon stiffness for maximising gastrocnemius efficiency during human walking and running. J. Theor. Biol. 252, 662–673.

Lussiana, T., Gindre, C., Hébert-Losier, K., Sagawa, Y., Gimenez, P. and Mourot, L. (2017). Similar running economy with different running patterns along the aerial-terrestrial continuum. Int. J. Sports Physiol. Perform. 12, 481–489.

Mercer, J. A. and Horsch, S. (2015). Heel-toe running: A new look at the influence of foot strike pattern on impact force. J. Exerc. Sci. Fit. 13, 29–34.

Miller, R. H. (2014). A comparison of muscle energy models for simulating human walking in three dimensions. J. Biomech. 47, 1373–1381.

Ogueta-Alday, A., Rodríguez-Marroyo, J. A. and García-López, J. (2014). Rearfoot striking runners are more economical than midfoot strikers. Med. Sci. Sports Exerc. 46, 580–585.

Patterson, M. A. and Rao, A. V. (2014). GPOPS-II. ACM Trans. Math. Softw. 41, 1–37.

Perl, D. P., Daoud, A. I. and Lieberman, D. E. (2012). Effects of footwear and strike type on running economy. Med. Sci. Sports Exerc. 44, 1335–1343.

Roper, J. L., Doerfler, D., Kravitz, L., Dufek, J. S. and Mermier, C. (2017). Gait Retraining from Rearfoot Strike to Forefoot Strike does not change Running Economy. Int. J. Sports Med. 38, 1076–1082.

Schache, A. G., Brown, N. A. T. and Pandy, M. G. (2015). Modulation of work and power by the human lower-limb joints with increasing steady-state locomotion speed. J. Exp. Biol. 218, 2472–2481.

Stearne, S. M., Alderson, J. A., Green, B. A., Donnelly, C. J. and Rubenson, J. (2014). Joint kinetics in rearfoot versus forefoot running: Implications of switching technique. Med. Sci. Sports Exerc. 46, 1578–1587.

Stearne, S. M., McDonald, K. A., Alderson, J. A., North, I., Oxnard, C. E. and Rubenson, J. (2016). The Foot’s Arch and the Energetics of Human Locomotion. Sci. Rep. 6,.

Swinnen, W., Hoogkamer, W., Delabastita, T., Aeles, J., De Groote, F. and Vanwanseele, B. (2019). Effect of habitual foot-strike pattern on the gastrocnemius medialis muscle-tendon interaction and muscle force production during running. J. Appl. Physiol. 126, 708–716.

Uchida, T. K., Hicks, J. L., Dembia, C. L. and Delp, S. L. (2016). Stretching your energetic budget: How tendon compliance affects the metabolic cost of running. PLoS One 11, 1–19.

Umberger, B. R. (2010). Stance and swing phase costs in human walking. J. R. Soc. Interface 7, 1329–40.

Umberger, B. R., Gerritsen, K. G. M. and Martin, P. E. (2003). A Model of Human Muscle Energy Expenditure. Comput. Methods Biomech. Biomed. Engin. 6, 99–111.

van der Zee, T. J., Lemaire, K. K. and van Soest, A. J. (2019). The metabolic cost of in vivo constant muscle force production at zero net mechanical work. J. Exp. Biol. 222, jeb199158.

Wächter, A. and Biegler, L. T. (2006). On the implementation of an interior-point filter line-search algorithm for large-scale nonlinear programming. Math. Program. 106, 25–57.

Waters, R. L. and Mulroy, S. (1999). The energy expenditure of normal and pathologic gait. Gait Posture 9, 207–231.

Zajac, F. E. (1989). Muscle and tendon: properties, models, scaling, and application to biomechanics and motor control. Crit. Rev. Biomed. Eng. 17, 359–410.

