## Supplementary figures for "Habitual foot strike pattern does not affect simulated Triceps Surae muscle metabolic energy consumption during running"

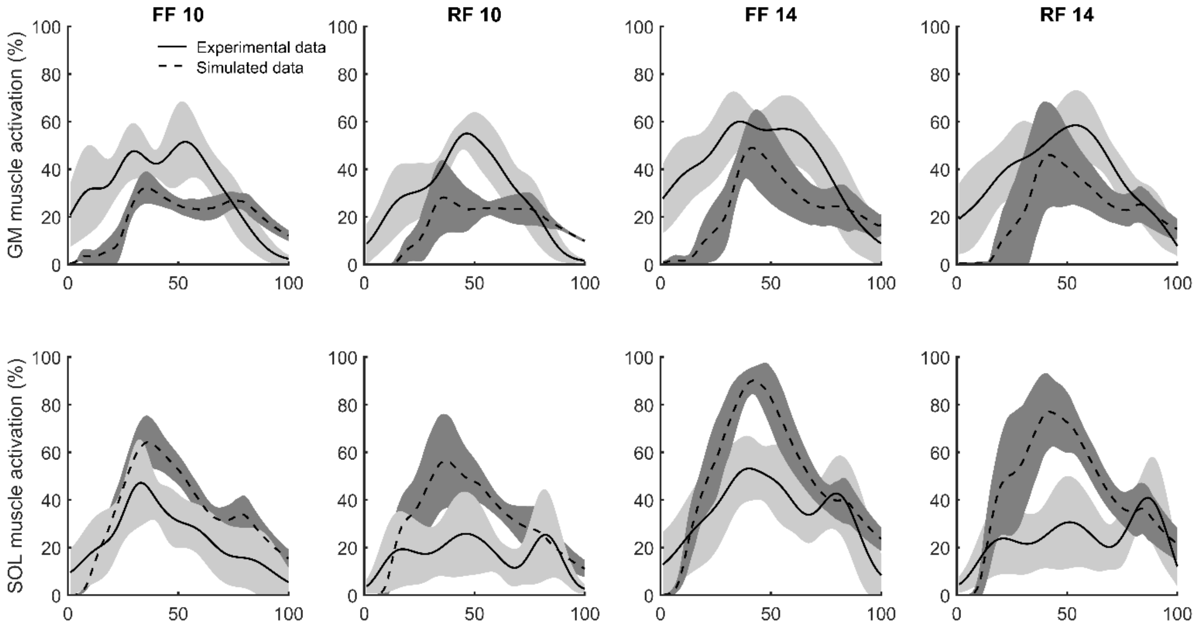


*Fig. S1. Gastrocnemius medialis (GM) and Soleus (SOL) experimental EMG data (solid) and simulated activation (dashed) in mid-/forefoot strikers (FF; GM: n = 9; SOL n = 8) and rearfoot strikers (RF; GM = 9; SOL = 8). Shaded area represent standard deviation.*

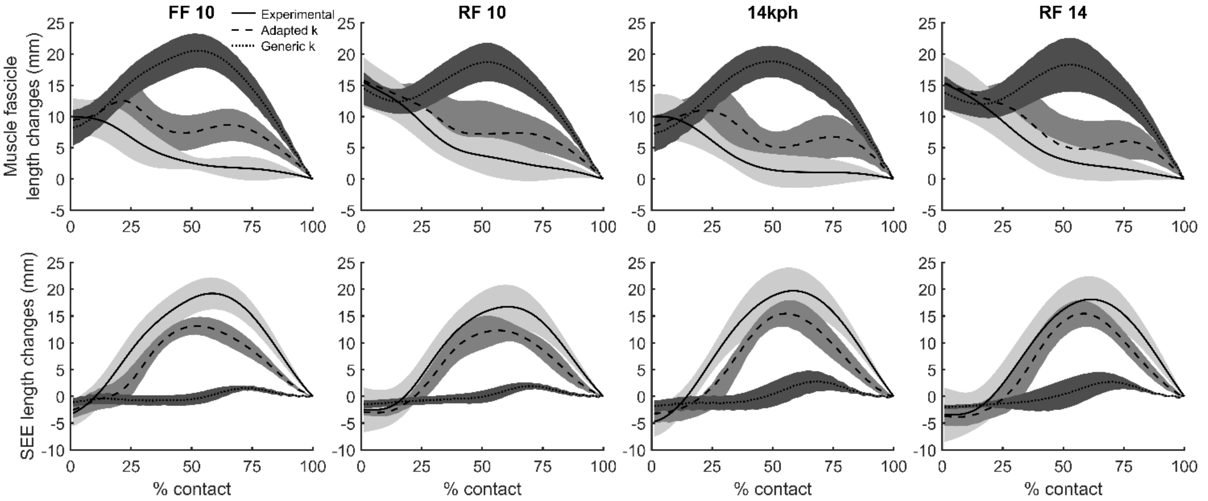


*Fig. S2. GM muscle fascicle and tendinous tissue (SEE) length changes during running in mid-/forefoot strikers (FF; n = 10) and rearfoot strikers (RF; n = 9). Experimental ultrasound data (solid), adapted simulation data (dashed) and generic simulation data (dotted).*

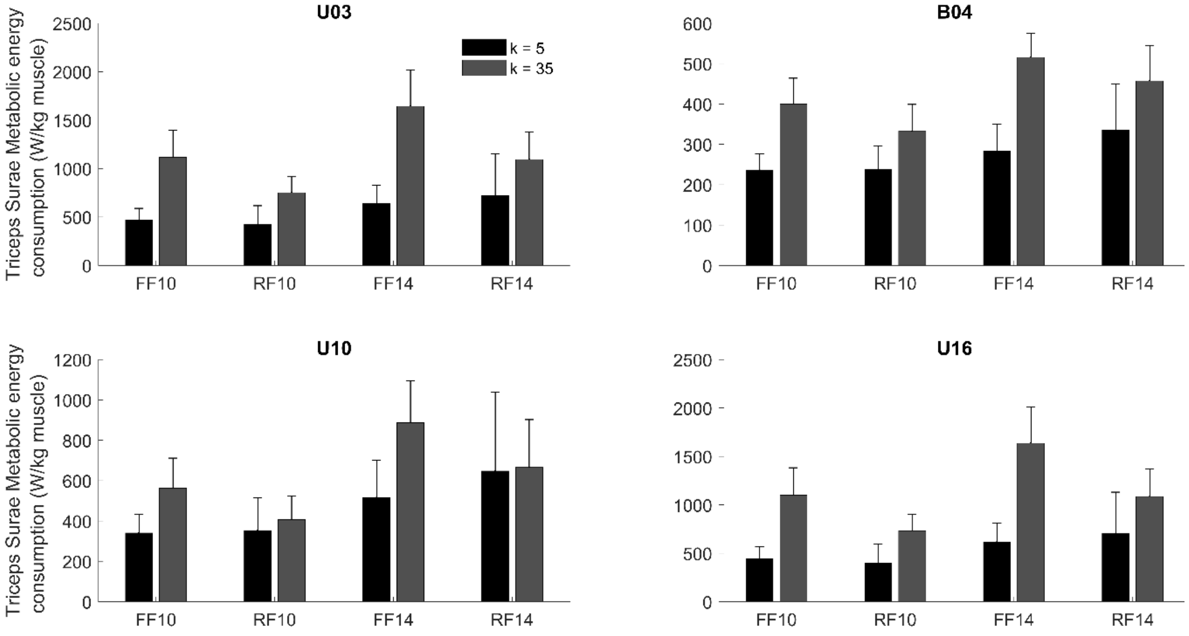


*Fig. S3. Estimated Triceps Surae muscle metabolic energy consumption when an adapted normalized Achilles tendon stiffness was used (k = 5, black) and when the generic normalized tendon stiffness was used (k = 35; grey) in mid-/forefoot strikers (FF; n = 10) and rearfoot strikers (RF; n = 9) at 10 and 14 km/h.*

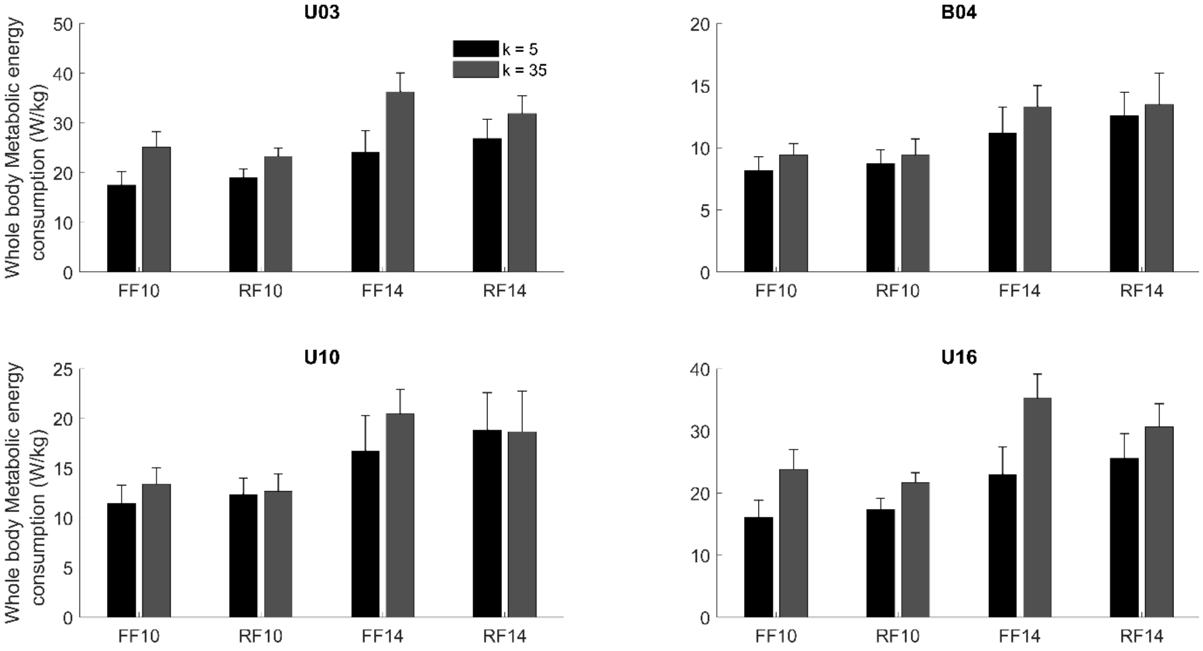


*Fig. S4. Estimated whole body metabolic energy consumption when an adapted normalized Achilles tendon stiffness was used (k = 5, black) and when the generic normalized tendon stiffness was used (k = 35; grey) in mid-/forefoot strikers (FF; n = 10) and rearfoot strikers (RF; n = 9) at 10 and 14 km/h.*
